# Central amygdala CRF pathways in alcohol dependence

**DOI:** 10.1101/134759

**Authors:** Giordano de Guglielmo, Marsida Kallupi, Matthew B. Pomrenze, Elena Crawford, Sierra Simpson, Paul Schweitzer, George F. Koob, Robert O. Messing, Olivier George

## Abstract

Alcohol withdrawal activates a neuronal ensemble in the central nucleus of the amygdala (CeA) that is responsible for high levels of uncontrolled alcohol drinking. However, the neuronal phenotypes and circuits controlled by these neurons are unknown. We investigated the cellular identity of this CeA neuronal ensemble and found that most neurons expressed corticotropin-releasing factor (CRF). Using *Crh*-Cre transgenic rats combined with *in vivo* optogenetics, we tested the role of CeA CRF neurons and their projections in excessive alcohol self-administration during withdrawal. Rats were injected with AAV-DIO-NpHR-eYFP or AAV-DIO-eYFP and implanted with optical fibers over the CeA. Animals were then exposed to chronic intermittent ethanol vapor to induce alcohol dependence. Inactivation of CeA CRF neurons decreased alcohol drinking in dependent rats to non-dependent levels and completely suppressed activation of the CeA neuronal ensemble (Fos^+^ neurons) during withdrawal. No effects were observed on water or saccharin self-administration. In a second experiment, CeA CRF neurons were infected with AAV-DIO-NpHR-eYFP and optical fibers were implanted into downstream projection regions, including the bed nucleus of the stria terminalis (BNST), lateral hypothalamus (LH), parasubthalamic nucleus (pSTN), substantia innominata (SI), and parabrachial nuclei (PBN). Optogenetic inactivation of CRF terminals in the BNST reduced alcohol drinking and withdrawal signs, whereas inactivation of all other projections had no effect. These results demonstrate that CeA CRF neurons and their projections to the BNST drive excessive alcohol drinking and withdrawal in dependent rats.

## Introduction

Alcoholism is a chronic relapsing disorder associated with compulsive drinking, loss of control over intake, and emergence of a negative emotional state during abstinence from the drug (Koob *et al*, 2004). Animal and human studies implicated the central nucleus of the amygdala (CeA) in alcohol use disorders (Janak and Tye, 2015). In fact, it has been shown that chronic alcohol use alters neuronal transmission in the CeA (Roberto *et al*, 2012) and that the CeA stores memory traces related to the smell and taste of alcohol that drives relapse (Barak *et al*, 2013). Importantly, activation of a specific neuronal ensemble in the CeA during alcohol withdrawal is causally related to the excessive alcohol drinking that is observed in alcohol-dependent rats (de Guglielmo *et al*, 2016). However, the cellular phenotypes of these neurons and brain regions that are controlled by this neuronal ensemble are unknown. Moreover, activation of corticotropin-releasing factor 1 (CRF_1_) receptors in the CeA is required for excessive alcohol drinking in dependent rats, but no direct evidence has been provided to date that activation of CeA CRF neurons is required for alcohol drinking. Indeed, extracellular levels of CRF in the CeA increase during withdrawal from chronic alcohol exposure (Koob, 2008; Koob *et al*, 2004; Merlo Pich *et al*, 1995; Olive *et al*, 2002), and systemic and intra-CeA administration of specific CRF_1_ antagonists reduce both the negative emotional states of alcohol withdrawal and alcohol drinking in dependent rats (Funk *et al*, 2006; Funk *et al*, 2007; Gehlert *et al*, 2007; Koob, 2008). Despite this substantial pharmacology evidence, no study has shown that the inactivation of CeA CRF neurons *per se* (not CRF_1_ receptors) is responsible for excessive alcohol drinking during acute withdrawal. This is critical because activation of CeA CRF_1_ receptors could result from the activation of CRF neurons that are located in other brain regions (bed nucleus of the stria terminalis [BNST], lateral hypothalamus [LH], parasubthalamic nucleus [pSTN]) that project to the CeA.

We first hypothesized that CRF neurons are critical for recruitment of the CeA neuronal ensemble during alcohol withdrawal. To test this hypothesis, we used *Crh*-Cre transgenic rats combined with *in vivo* optogenetics and immediate early gene brain mapping. We then dissected the role of the different downstream CeA CRF pathways in alcohol dependence using optogenetic inactivation of CeA CRF terminals that target the BNST (CRF^CeA-BNST^), LH and pSTN (CRF^CeA-LH/pSTN^), substantia innominata (SI; CRF^CeA-SI^), and parabrachial nucleus (PBN; CRF^CeA-PBN^; Pomrenze *et al*, 2015).

## Materials and Methods

### Subjects

Adult male *Crh*-Cre rats, weighing 200-225 g at the beginning of the experiments, were housed in groups of two per cage (self-administration groups) in a temperature-controlled (22°C) vivarium on a 12 h/12 h light/dark cycle (lights on at 10:00 PM) with *ad libitum* access to food and water. All of the behavioral tests were conducted during the dark phase of the light/dark cycle. All of the procedures adhered to the National Institutes of Health Guide for the Care and Use of Laboratory Animals and were approved by the Institutional Animal Care and Use Committee of The Scripps Research Institute.

### Operant self-administration

Self-administration sessions were conducted in standard operant conditioning chambers (Med Associates, St. Albans, VT, USA). For alcohol self-administration studies, animals were first trained to self-administer 10% (w/v) alcohol and water solutions until a stable response pattern (20 ± 5 rewards) was maintained. Rats were then subjected to an overnight session in the operant chambers with access to one lever (right lever) that delivered water (fixed-ratio 1 [FR1]). Food was available *ad libitum* during this training. After 1 day off, rats were subjected to a 2 h session (FR1) for 1 day and a 1 h session (FR1) the next day, with one lever delivering alcohol (right lever). All of the subsequent alcohol self-administration sessions lasted 30 min. Rats were allowed to self-administer a 10% (w/v) alcohol solution (right lever) and water (left lever) on an FR1 schedule of reinforcement (i.e., each operant response was reinforced with 0.1 ml of the solution). For saccharin self-administration, rats underwent daily 30 min FR1 sessions. Responses on the right lever resulted in the delivery of 0.1 ml of saccharin (0.04%, w/v). Lever presses on the left lever delivered 0.1 ml of water. This procedure lasted 13 days until a stable baseline of intake was achieved.

### Alcohol vapor chambers

Rats were trained to self-administer alcohol in the operant chambers. Once a stable baseline of alcohol intake was reached, the rats were made dependent by chronic intermittent exposure to alcohol vapors. They underwent cycles of 14 h on (blood alcohol levels during vapor exposure ranged between 150 and 250 mg%) and 10 h off, during which behavioral testing for acute withdrawal occurred (i.e., 6-8 h after vapor was turned off, when brain and blood alcohol levels are negligible). In this model, rats exhibit somatic and motivational signs of withdrawal (Vendruscolo and Roberts, 2014).

### Operant self-administration and withdrawal scores during alcohol vapor exposure

Behavioral testing occurred three times per week. Rats were tested for alcohol or saccharin (and water) self-administration on an FR1 schedule of reinforcement in 30-min sessions. Behavioral signs of withdrawal were measured using a rating scale that was adapted from a previous study (Macey *et al*, 1996) and included ventromedial limb retraction (VLR), irritability to touch (vocalization), tail rigidity, abnormal gait, and body tremors. Each sign was given a score of 0-2, based on the following severity scale: 0 = no sign, 1 = moderate, 2 = severe. The sum of the four observation scores (0-8) was used as an operational measure of withdrawal severity.

### Intracranial surgery

Rats were anesthetized with isoflurane (5% v/v) and secured in a stereotaxic frame. Cre-dependent adeno-associated viruses AAV-DIO-eNpHR3.0-eYFP or AAV-DIO-eYFP (UNC Vector Core, University of North Carolina, Chapel Hill) were injected bilaterally or unilaterally in the CeA (coordinates from Bregma: −2.6 mm AP, ±4.2 mm ML, −6.1 mm DV from skull surface) through a stainless-steel injector that was 2 mm longer than the guide cannula (so that the tip protruded into the area) and connected to a 10 µl Hamilton syringe, which was controlled by an UltraMicroPump (WPI, Sarasota, FL, USA). Virus was injected (1.0 µl, 100 nl/min) over 10 min followed by an additional 10 min to allow the diffusion of viral particles. Rats used for behavioral experiments were then implanted bilaterally or unilaterally with chronic fiber optics above the CeA (coordinates from Bregma: AP −2.6 mm, ML ± 4.2 mm, DV −8.1 mm from skull surface), or unilaterally above the BNST (AP −0.1, ML ± 1.4, DV −6.7 from skull surface), the LH/pSTN (AP −4.2, ML ± 1.2, DV −8.7 from skull surface), the SI (AP −0.1, ML ± 1.8, DV −8.5 from skull surface) or the PBN (AP −9.2, ML ± 1.8, DV −6.5 from skull surface). Following surgery, rats were allowed to recover for 1 week.

### Locomotor activity

Locomotor activity and anxiety-like behavior were evaluated in an opaque open field apparatus (100 cm × 100 cm × 40 cm) that was divided into 20 cm × 20 cm squares. On the experimental day, the animals were placed in the center of the open field, and locomotor activity (number of lines crossed), the time in the center, total distance traveled, and entries into the center were scored for 10 min. Behavior was videotaped and analyzed with Any Maze software.

### Optical inhibition

Optical probes were constructed, in which the fiber optic (200 µm core, multimode, 0.37 NA) was inserted and glued into a ceramic ferrule. The fiber optic and ferrule were then glued to a metallic cannula and closed with a dust cap. The fiber optic length was then adjusted based on the brain area of interest to not leave a gap between the skull and cannula during surgery. The other end of the fiber optic (FC/PC connection) was attached to a fiber splitter (2×1) that permitted simultaneous, bilateral illumination. The single end of the splitter was attached to a rotating optical commutator (Doric Lenses) to permit free movement of the rat. The commutator was connected to a fiber that was connected to a laser (DPSS, 200 or 300 mW, 532 nm, with a multimode fiber coupler for an FC/PC connection; OEM Laser Systems). Prior to the experiments, the light output of the fiber optic was adjusted to approximately 10 mW, as measured by an optical power meter. Based on measurements in the mammalian brain (Deisseroth, 2012), assuming a geometric loss of light, a light output of 10 mW that is measured by a standard optical power at the tip of a fiber with an NA of 0.37 and fiber core radius of 200 µm will produce ~1 mW/mm^2^ of light up to 1 mm directly away from the fiber tip, which is the minimum amount necessary to produce opsin activation (Gradinaru *et al*, 2009; Tye *et al*, 2011). Based on *in vivo* measurements of the shape of the light output in mammalian brain tissue, these parameters would be expected to provide sufficient light for opsin activation in at least 0.4 mm^3^ of tissue (Yizhar *et al*, 2011). Fiber optics were implanted at the same time as the virus injections, and the animals were habituated to tethering to the commutator for several self-administration sessions before beginning the experiments. On the test days, the rats received optical inhibition paired with alcohol self-administration sessions, locomotor activity assessments, or withdrawal score assessments.

### Recruitment of CeA CRF neurons during alcohol withdrawal

Rats were trained to self-administer 10% alcohol and made dependent on alcohol using the chronic intermittent exposure model. After the stabilization of excessive drinking, the animals were transcardially perfused. The brains were then harvested, postfixed, and sectioned at 40 µm.

### Effect of optogenetic inhibition of CeA CRF neurons on alcohol and saccharin self-administration in nondependent and alcohol-dependent rats

Animals were injected with AAV-DIO-eNpHR3.0-eYFP or AAV-DIO-eYFP and implanted with bilateral fiber optics over the CeA. One week after surgery, the rats were trained to self-administer 10% alcohol until they reached a stable baseline of intake. At this point, rats were tested for alcohol self-administration during optical inhibition of CeA CRF neurons. Sham optical inhibition sessions were performed before and after the test session. The animals were then placed in the alcohol vapor chambers and 3 weeks later underwent the alcohol self-administration escalation phase. When the escalation of alcohol intake was achieved, animals were tested for the effects of the optical inhibition of CeA CRF neurons as described above. The effect of optical inhibition on somatic withdrawal signs was also assessed.

At the end of this phase, rats were trained to self-administer saccharin for several sessions until a stable baseline of responses was reached. They were then tested for the effects of the optical inhibition of CeA CRF neurons on saccharin self-administration. The effects of optical inhibition on locomotor activity were also assessed.

### Dissection of the role of different CeA CRF pathways on alcohol and saccharin self-administration in alcohol-dependent rats

*Crh*-Cre rats were bilaterally infused with AAV-DIO-NpHR-eYFP in the CeA, unilaterally implanted with fiber optics over the CeA, BNST, PBN, SI, or LH, and made dependent on alcohol using the chronic intermittent exposure model. After escalation of drinking, the animals were tested for the effect of the optogenetic inhibition of CeA CRF projections on alcohol drinking, alcohol withdrawal signs, and saccharin self-administration.

### Immunohistochemistry

*Crh*-Cre rats were unilaterally prepared with AAV-DIO-NpHR-EYFP and an optic fiber in the CeA and tested for the effect of inhibition of CeA CRF neurons for 30 min during withdrawal (8 h) from chronic intermittent alcohol. Ninety minutes later, the animals were deeply anesthetized and perfused with 100 mL of phosphate-buffered saline (PBS) followed by 400 mL of 4% paraformaldehyde. Brains were postfixed in 4% paraformaldehyde overnight and transferred to 30% sucrose in PBS/0.1% azide solution at 4°C for 2-3 days. Brains were frozen in powdered dry ice and sectioned on a cryostat. Coronal sections were cut 40 µm thick between bregma + 4.2 and −6.48 mm (Paxinos and Watson, 2005) and collected free-floating in PBS/0.1% azide solution. Following three washes in PBS, the sections were incubated in 1% hydrogen peroxide/PBS to quench endogenous peroxidase activity, rinsed three times in PBS, and blocked for 60 min in PBS that contained 0.3% TritonX-100, 1 mg/ml bovine serum albumin, and 5% normal donkey serum. Sections were incubated for 24 h at 4°C with rabbit monoclonal anti-Fos antibody (Cell Signaling Technologies, catalog no. 2250) diluted 1:1000 in PBS/0.5% Tween20 and 5% normal donkey serum. The sections were washed again with PBS and incubated for 1 h in undiluted Rabbit ImmPress HRP reagent (Vector Laboratories). After washing in PBS, the sections were developed for 2-6 min in Vector peroxidase DAB substrate (Vector Labs) enhanced with nickel chloride. Following PBS rinses, the sections were mounted on coated slides (Fisher Super Frost Plus), air dried, dehydrated through a graded series of alcohol, cleared with Citrasolv (Fisher Scientific), and coverslipped with DPX (Sigma).

Quantitative analysis to obtain unbiased estimates of the total number of Fos+ cell bodies was performed on a Zeiss Axiophot Microscope equipped with MicroBrightField Stereo Investigator software (Colchester, VT, USA), a three-axis Mac 5000 motorized stage (Ludl Electronics Products, Hawthorne, NY, USA), a Q Imaging Retiga 2000R color digital camera, and PCI color frame grabber.

Fluorescent immunohistochemistry was used to characterize the type (CRF) and number (NeuN) of neurons that were activated (Fos^+^) during alcohol withdrawal. Coronal sections were cut 40 µm thick between bregma −2.0 and −5.0mm (Paxinos *et al*, 2005). We determined the proportion of all neurons expressing Fos during alcohol withdrawal by double-labeling for Fos and the neuron-specific protein NeuN, as well as the population of activated CRF neurons in the CeA by double labeling for Fos and CRF.

For Fos/NeuN and Fos/CRF double-labeling, 40 µm sections were washed three times in PBS and permeabilized/blocked for 60 min in PBS with 0.2% Triton X-100, 5% normal donkey serum, and 3% bovine serum albumin (blocking solution). Sections were incubated in primary antibodies diluted in blocking solution for 24 h on a shaker at room temperature. Primary antibodies were used at the following concentrations: anti-Fos (Millipore, catalog no. AB4532, 1:500) and anti-NeuN (Millipore, catalog no. MAB377, 1:1000). Sections were washed three times for 10 min in PBS and incubated with fluorescently labeled secondary antibodies diluted in PBS for 2 h on a shaker at room temperature. Species-specific secondary antibodies used were Alexa Fluor 488 (1:500, Invitrogen, catalog no. A-10042), Alexa Fluor 568 (1:500, Invitrogen, catalog no. S-11249), and Alexa Fluor 647 (1:500, Invitrogen, catalog no. A-21447). After labeling, the sections were washed three times in PBS for 10 min, mounted on Fisher Superfrost Plus slides (catalog no. 12-550-15), and coverslipped with PVA-Dabco (Sigma).

Three sections were bilaterally analyzed for each rat. Cells were identified as neurons based on standard morphology, and only neurons with a focused nucleus within the boundary of the CeA were counted. Counts from all images for each rat were averaged so that each rat was an *n* of 1.

### Confocal acquisition and three-dimensional analysis

Three-dimensional stacks of images were acquired with a 780 Laser Scanning Confocal microscope (Zeiss) using a 20× (1 µm image slice), 40× (0.6 µm image slice), or 63× (0.2 µm image slice) objective to observe the entirety of the CeA. The system is equipped with a stitching stage and Zen software to reintegrate the tiled image stacks. Stitched z-series images of the entire CeA were imported into Imaris software (Bitplane-Andor) and FIJI for quantification.

### Slice preparation for whole-cell recordings

*Crh*-Cre rats (*n* = 5) expressing AAV-DIO-eNpHR-EYFP in the CeA for 4-5 weeks were deeply anesthetized with 3% isoflurane and then transcardially perfused with ice-cold oxygenated sucrose solution. The rats were then decapitated, and the brains were rapidly removed and placed into oxygenated (95% O_2_, 5% CO_2_) ice-cold cutting solution that contained 206 mM sucrose, 2.5 mM KCl, 1.2 mM NaH_2_PO_4_, 7 mM MgCl_2_, 0.5 mM CaCl_2_, 26 mM NaHCO_3_, 5 mM glucose, and 5 mM HEPES. CeA slices (300 um) thick were cut on a Vibratome (Leica VT1000S, Leica Microsystems, Buffalo Grove, IL, USA) and transferred to oxygenated artificial cerebrospinal fluid (aCSF) that contained 130 mM NaCl, 2.5 mM KCl, 1.25 mM NaH_2_PO_4_, 1.5 mM MgSO_4_·7H_2_O, 2.0 mM CaCl_2_, 24 mM NaHCO_3_, and 10 mM glucose. Slices were first incubated for 30 min at 37°C and then kept at room temperature for the remainder of the experiment. Individual slices were transferred to a recording chamber that was mounted on the stage of an upright microscope (Olympus BX50WI, Waltham MA, USA). Recordings were performed in continuous oxygenated aCSF that was perfused at a rate of 2-3 ml/min. Neurons were visualized with a 60× water immersion objective (Olympus), infrared differential interference contrast optics, and a charge-couple device camera (EXi Blue, QImaging, Surrey, Canada). Whole-cell recordings were performed using current clamp mode with a Multiclamp 700B amplifier, Digidata 1440A, and pClamp 10 software (Molecular Devices, Sunnyvale, CA, USA). Patch pipettes (4-7 MΏ) were pulled from borosilicate glass (Warner Instruments, Hamden, CT, USA) and filled with the following internal solution: 70 mM KMeSO_4_, 55 mM KCl, 10 mM NaCl, 2 mM MgCl_2_, 10 mM HEPES, 2 mM Mg-ATP, and 0.2 mM Na-GTP.

To study light-induced neuronal inhibition, a green laser (532 nm) was switched on for 6.5 s by a Master-8 stimulator (AMPI, Jerusalem, Israel) to generate trains at a frequency of 15 Hz. Neurons that expressed eNpHR within the CeA were visualized with differential interference contrast and widefield fluorescence imaging using an Olympus 60× immersion objective. To identify eNpHR-expressing neurons, a Lambda DG-4 light source was used with an in-line EX540/EM630 filter set. A Mosaic 3 pattern illuminator (Andor Instruments, Belfast, UK) coupled to a 532 nm light-emitting diode (CoolLED Limited, Andover, UK) was attached to the microscope and used for light delivery (10 mW/mm^2^, to approximate output in the behavioral experiments) through the objective within the slice preparation.

### Histology

Following completion of behavioral experiments, the rats were deeply anaesthetized and transcardially perfused. Brains were removed, postfixed, cryoprotexted in 30% sucrose, and sectioned at 40 µm into coronal sections containing the CeA, BNST, and PBN. Brain slices were mounted on microscope slides and expression of viral vectors and/or optical fiber placements were examined for all of the rats using fluorescent microscopy. Rats that had no eYFP expression in the CeA or had fiber placements outside the target regions of interest were excluded from the behavioral analysis.

### Statistical analysis

Data are expressed as mean ± SEM. For comparisons between only two groups, *p* values were calculated using paired or unpaired *t*-tests as described in the figure legends. Comparisons across more than two groups were made using one-way analysis of variance (ANOVA), and two-way ANOVA was used when there was more than one independent variable. The Newman Keuls *post hoc* test was used following significance in the ANOVA. Withdrawal signs were analyzed by the nonparametric Mann-Whitney *U* statistic, followed by Dunn’s multiple-comparison test. The standard error of the mean is indicated by error bars for each group of data. Differences were considered significant at *p* < 0.05. All of the data were analyzed using Statistica 7 software.

## Results

### CeA CRF neurons are recruited during alcohol withdrawal

At the end of the escalation phase and exposure to chronic intermittent ethanol vapor, rats averaged 45.1 ± 7.2 responses for alcohol compared with 17.5 ± 3.4 responses before exposure to chronic intermittent ethanol (*p* < 0.01; Fig 1.A). After the stabilization of excessive drinking, the animals were transcardially perfused. Withdrawal from alcohol vapor produced a significant increase in Fos^+^ neurons in the CeA (*t*_8_ = 6.16, *p* < 0.001; Fig. 1B). This recruitment was limited to a small subpopulation of neurons in the CeA. Double-labeling with Fos/NeuN revealed that only 7-8% of all CeA neurons (NeuN+) were also Fos^+^ (*t*_8_ = 22.4, *p* < 0.01; Supplementary Fig. S1). Double Fos-CRF immunohistochemistry showed that the total number of CRF neurons did not change between withdrawal rats and naive rats (*t*_8_ = 0.58, *p* = 0.05; Fig. 1C), but the number of Fos^+^/CRF^+^ neurons in the CeA significantly increased during withdrawal (*t*_8_ = 6.24, *p* < 0.001; Fig. 1D) and represented the majority of Fos^+^ neurons (80%).

**Figure 1.**
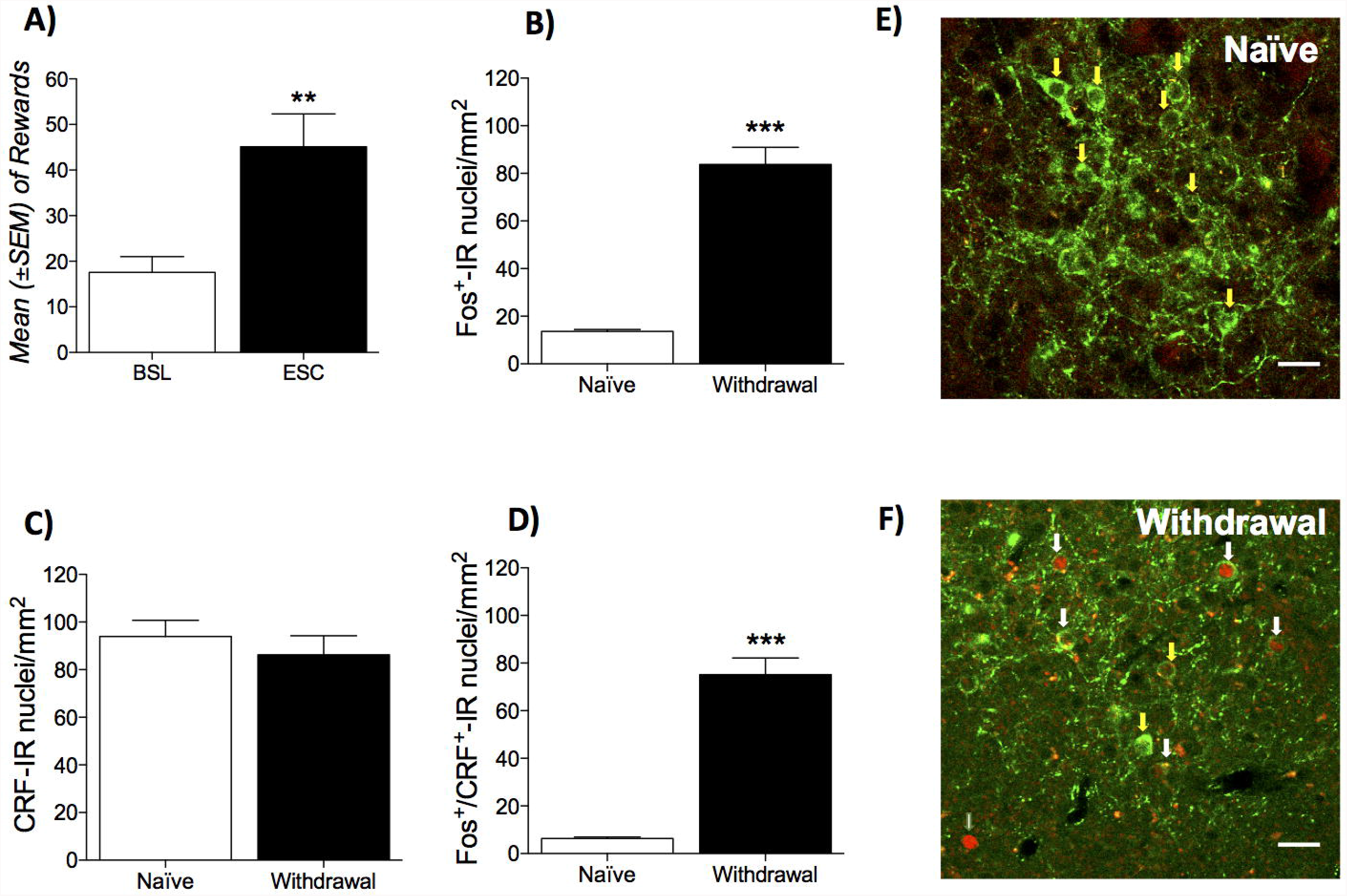
Alcohol withdrawal recruits CRF neurons in the CEA. (**A**) Escalation of alcohol drinking after chronic intermittent alcohol exposure. \*\**p* < 0.01, *vs*. baseline. (**B**) Number of Fos+ nuclei per mm^2^ in the CeA. (**C**) Double Fos-CRF immunohistochemistry showed that the total number of CRF neurons did not change between withdrawal rats and naive rats, but (**D**) the number of Fos+/CRF+ neurons in the CeA dramatically increased during withdrawal. (**E, F**) Representative pictures of double Fos-CRF immunohistochemistry in the CeA in naive (**E**) and alcohol withdrawal (**F**) rats. The data are expressed as mean ± SEM. \*\**p* < 0.01, \*\*\**p* < 0.001, *vs*. naive.

### Validation of NpHR in CeA CRF neurons

To confirm the functionality of NpHR, electrophysiological recordings of neuronal activity from single neurons were performed in acute CeA slices from *Crh*-Cre rats. CeA neurons were depolarized by current injection to evoke the sustained firing of action potentials, and CRF^+^ neurons were identified by visualization of eYFP fluorescence (Zhao *et al*, 2011). In six neurons that showed fluorescence that were held at −52.9 ± 0.4 mV, activation of a green laser (532 nm) using a 6.5 s train at 15 Hz (Haubensak *et al*, 2010) elicited hyperpolarization of 6.5 ± 1.0 mV, concomitantly suppressing the firing of action potentials (Fig. 2C). In another six neurons that did not show fluorescence, similar exposure to the laser (6.5 s at 15 Hz) did not elicit any significant effect on resting potential (−51.8 ± 0.5 mV hold, 0.5 ± 0.3 mV hyperpolarization; Fig. 2D).

**Figure 2.**
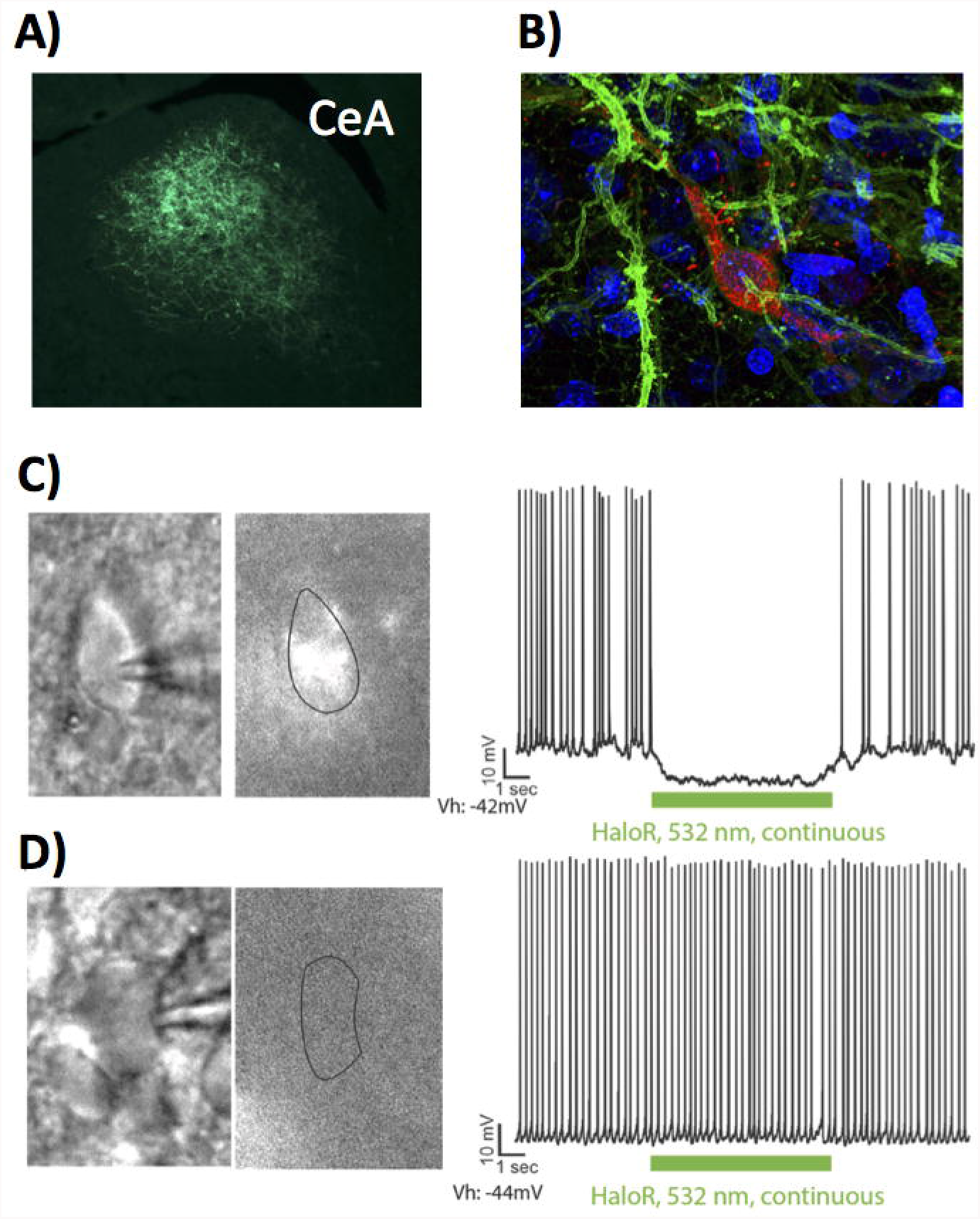
Validation of the *Crh*-Cre rat. (**A**) Cre-dependent eYFP expression in the CeA. (**B**) Cre-dependent eYFP colocalizes with CRF immunoreactivity in the CeA. (**C**) Infrared (*Left*) and YFP fluorescence (middle) images of a CeA neuron. (*Right*) Current clamp recording of the same neuron (held at −50 mV), depicting hyperpolarization in response to delivery of a 6.5 s train (asterisks) of green light. (**D**) In this CeA neuron (held at −52 mV) that did not present fluorescence, similar light stimulation did not elicit a response.

### Optogenetic inhibition of CeA CRF neurons reduces alcohol self-administration in alcohol-dependent rats

In non-dependent animals, inhibition of CeA CRF neurons had no effect in both NpHR and control rats. The ANOVA did not reveal a main effect of group (*F*_1,14_ = 1.30, *p* > 0.05) or treatment (*F*_1,28_ = 0.03, *p* > 0.05) or a group × treatment interaction (*F*_1,28_ = 2.10, *p* > 0.05). Water self-administration was unaffected by the green laser, with no effect of group (*F*_1,14_ = 0.17, *p* > 0.05) or treatment (*F*_1,28_ = 0.82, *p* > 0.05) and no group × treatment interaction (*F*_1,28_ = 1.07, *p* > 0.05; Fig. 3A).

**Figure 3.**
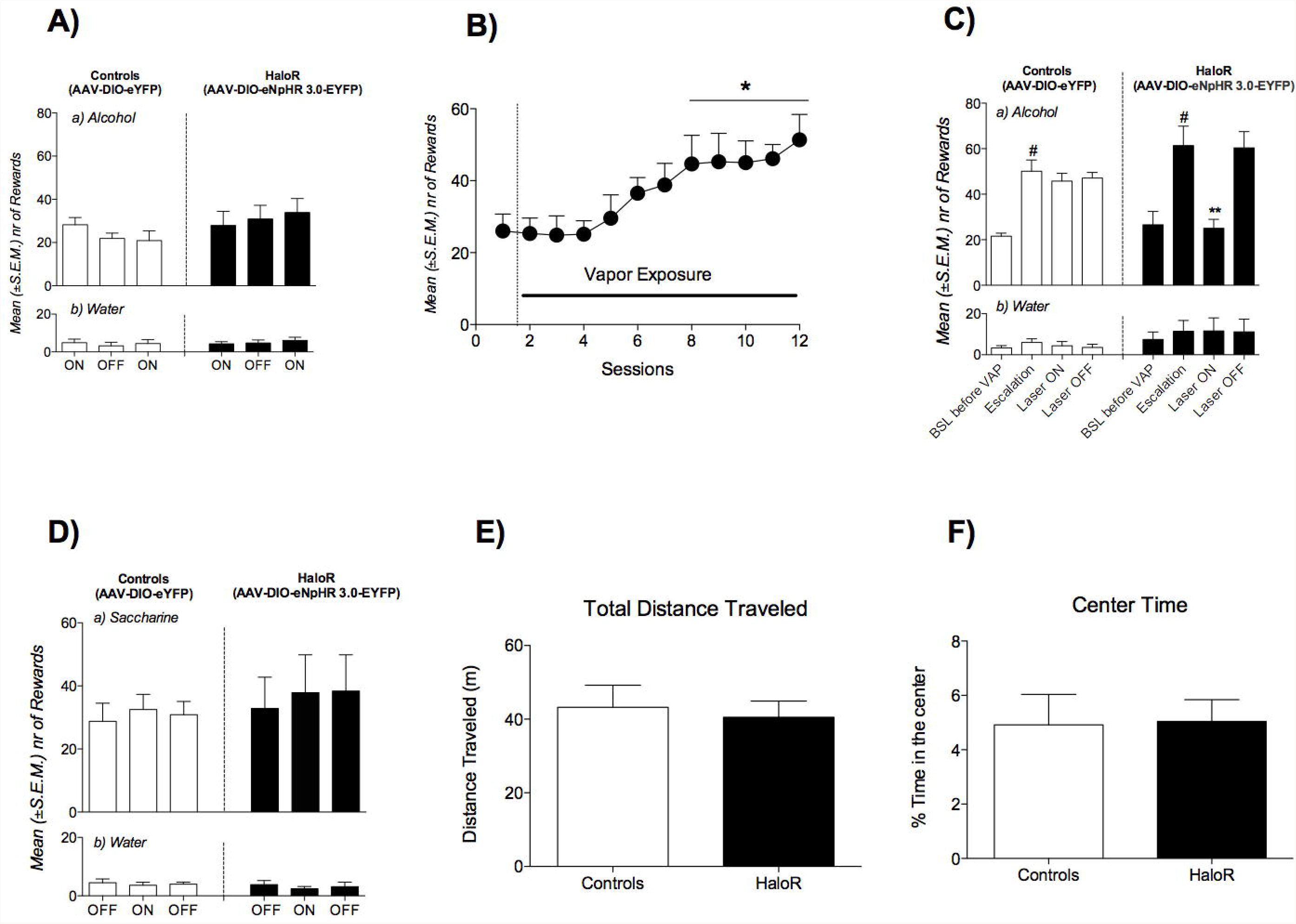
(**A**) Effect of optogenetic inhibition of CeA CRF neurons on alcohol (**a**) and water (**b**) self-administration in non-dependent rats. (**B**) Escalation of alcohol self-administration during alcohol vapor exposure. \**p* < 0.05, *vs*. baseline. (**C**) Effect of optogenetic inhibition of CeA CRF neurons on alcohol (**a**) and water (**b**) self-administration in alcohol-dependent rats. ^#^*p* < 0.05, vs. baseline (BSL); \*\**p* < 0.01, *vs*. Escalation. (**D**) Effect of optogenetic inhibition of CeA CRF neurons on saccharin (**a**) and water (**b**) self-administration. (**E, F**) Effect of optogenetic inhibition of CeA CRF neurons on total distance traveled (**E**) and time spent in the center (**F**) in the open field test. The data are expressed as mean ± SEM.

After 6 weeks of chronic intermittent alcohol vapor exposure, the animals escalated their alcohol consumption (*F*_1,28_ = 0.82, *p* < 0.001). The Newman Keuls *post hoc* test indicated a significant increase in alcohol self-administration starting from session 8 to session 12 compared with the first day of vapor exposure (*p* < 0.05; Fig. 3B). Fig. 3C shows the effects of the optogenetic inhibition of CeA CRF neurons on alcohol (Fig. 3C.a) and water (Fig. 3C.b) self-administration in alcohol-dependent rats. The mixed-factorial ANOVA indicated that both controls and NpHR rats escalated their alcohol intake after 5 weeks of vapor exposure (*F*_1,14_ = 14.9, *p* < 0.01; *p* < 0.05, *vs*. baseline pre-vapor, Newman Keuls *post hoc* test). Activation of the green laser selectively reduced alcohol self-administration in NpHR-injected rats. The ANOVA revealed a significant effect of treatment (*F*_2,28_ = 12.7, *p* < 0.001) and a group × treatment interaction (*F*_2,28_ = 8.38, *p* < 0.01) but no effect of group (*F*_1,14_ = 1.74, *p* > 0.05). The Newman Keuls *post hoc* test indicated a selective reduction of alcohol intake in NpHR-injected rats. No effect of the inhibition of CeA CRF neurons was detected on water self-administration, with no effect of group (*F*_1,14_ = 0.7, *p* > 0.05) or treatment (*F*_2,28_ = 0.22, *p* > 0.05) and no group × treatment interaction (*F*_2,28_ = 2.44, *p* > 0.05).

To verify that the effect of optical inhibition was not attributable to nonspecific effects of prolonged (30 min) illumination of the CeA with the green laser, we measured alcohol drinking after only 5 min of illumination. No difference in the magnitude of the effects was observed between the first 5 min and entire 30 min session (Supplementary Fig. S2A).

To further test the behavioral specificity of optical inhibition of CeA CRF neurons on alcohol drinking, the animals were trained to self-administer saccharin and tested for the effects of the inhibition of CeA CRF neurons on saccharin intake. The mixed-factorial ANOVA did not reveal an effect of activation of the green laser in controls or NpHR rats on saccharin intake (group: *F*_1,14_ = 0.27, *p* > 0.05; treatment: *F*_2,28_ = 0.48, *p* > 0.05; group × treatment interaction: *F*_2,28_ = 0.06, *p* > 0.05) or water intake (group: *F*_1,14_ = 0.57, *p* > 0.05; treatment: *F*_2,28_ = 1.13, *p* > 0.05; group × treatment interaction: *F*_2,28_ = 0.27, *p* > 0.05).

The animals were finally tested in the open field for effects on locomotor activity and anxiety-like behavior. The *t*-tests did not reveal significant effects of exposure to the green laser between controls and NpHR-injected rats on the total distance traveled (*t*_8_ = 0.36, *p* > 0.05) or time spent in the center (*t*_8_ = 0.36, *p* > 0.05). Therefore optogenetic silencing of CeA CRF neurons selectively reduces alcohol self-administration in dependent rats without affecting locomotor activity or anxiety.

### Unilateral inhibition of CeA CRF neurons reduces alcohol self-administration and Fos induction in the CeA

A separate group of *Crh*-Cre rats was bilaterally infused with AAV-DIO-NpHR-eYFP and implanted with unilateral fiber optics over the CeA (hemispheres with fiber optics were counterbalanced across al groups). The effect of the optogenetic inhibition of CeA CRF neurons on alcohol drinking, somatic signs of withdrawal, and saccharin self-administration were measured using the same chronic intermittent alcohol exposure protocol as in the previous experiments.

For alcohol drinking, the two-way repeated-measures ANOVA, with time and laser illumination as within-subjects factors, revealed significant effects of time (*F*_2,22_ = 16.36, *p* < 0.001) and laser illumination (*F*_1,11_ = 6.62, *p* < 0.05) and a time × laser illumination interaction (*F*_2,22_ = 13.15; *p* < 0.001). The Newman Keuls *post hoc* test showed a significant reduction of alcohol self-administration when the green laser was turned on (*p* < 0.01, *vs*. laser OFF; Fig. 4A).

**Figure 4.**
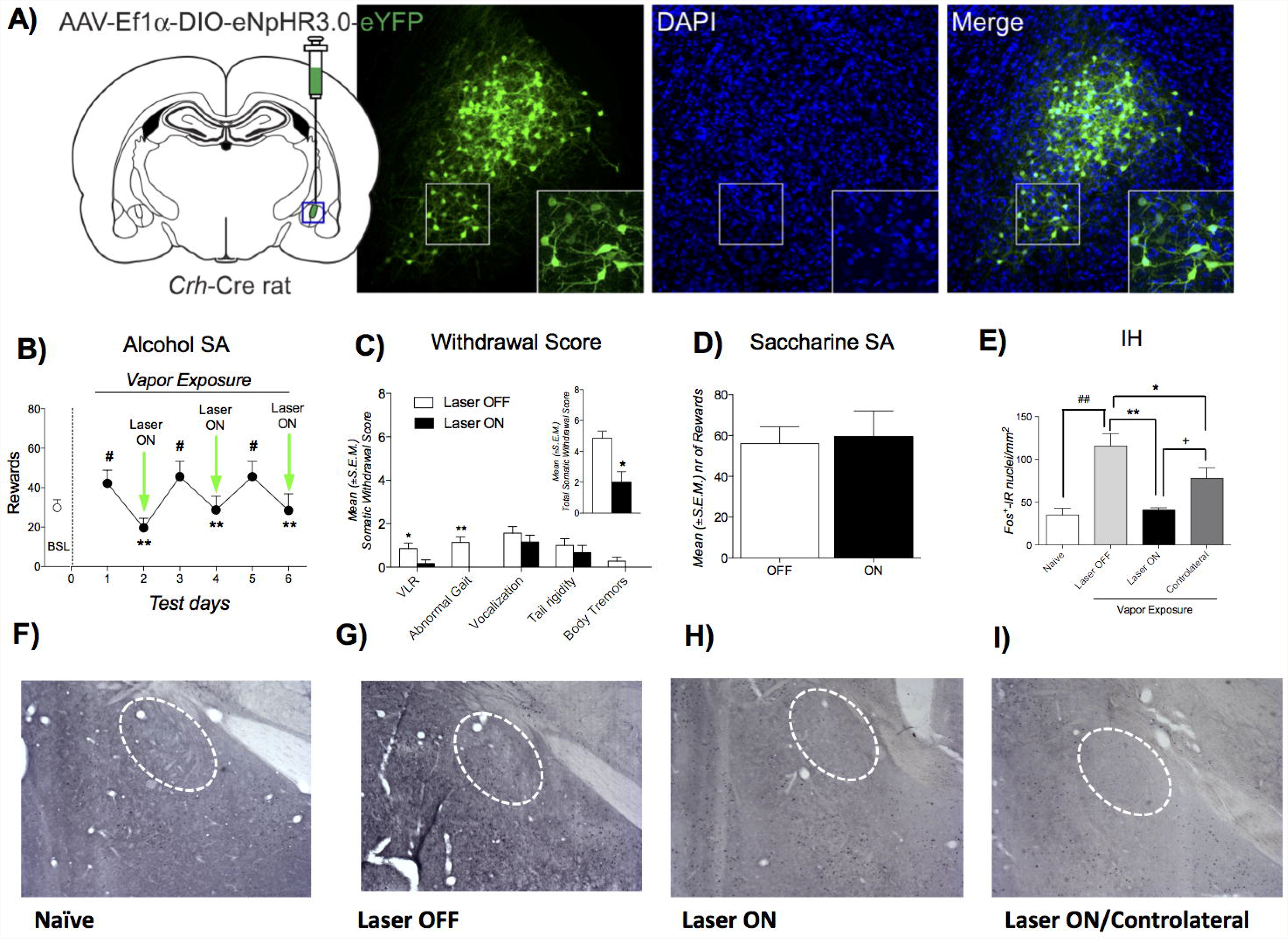
(**A**) Representative images of NpHR injection in the CeA. (**B**) Time-course of the effect of unilateral optogenetic inactivation of CeA CRF neurons. ^#^*p* < 0.05, *vs*. baseline (BSL); \*\**p* < 0.01, *vs*. laser OFF. (**C**) Effect of CeA CRF neuron inhibition on somatic withdrawal signs. \**p* < 0.05, vs. laser OFF. (**D**) Effect of unilateral optogenetic inhibition of CeA-CRF neurons on saccharin self-administration. (**E**) Effect of CeA CRF neuron inhibition for 30 min during withdrawal (8 h) from chronic intermittent alcohol exposure on CeA Fos immunoreactivity. ^##^*p* < 0.01, *vs*. naive; \**p* < 0.05, \*\**p* < 0.01, *vs*. laser OFF; ^+^*p* < 0.05, *vs*. contralateral. (**F-I)** Representative pictures of the different experimental conditions.

A separate one-way ANOVA indicated that the animals presented escalation of alcohol intake compared with baseline pre-vapor exposure (*F*_6,66_ = 9.23, *p* < 0.001). The escalation of alcohol drinking was observed only on days when the laser was off (*p* < 0.001) and was completely blocked by the optogenetic inhibition of CeA-CRF neurons, indicated by the Newman Keuls *post hoc* analysis.

Immediately before the last two alcohol self-administration sessions, rats were observed for somatic signs of withdrawal. As shown in Fig. 4B, the laser ON group exhibited significant decreases in ventromedial limb retraction (Mann-Whitney *U* = 9.00, *p* < 0.05) and abnormal gait (*U* = 3.000, *p* < 0.01). The sum of the five rating scores revealed a significant decrease in overall withdrawal severity (*U* = 4.5, *p* < 0.05; Fig. 4B, inset) during inhibition of CeA CRF neurons. The *t*-test did not indicate an effect of the inhibition of CeA CRF neurons on saccharin self-administration (*t*_11_ = 0.321, *p* > 0.05; Fig. 4C).

A separate group of rats was tested for the effect of inhibition of CeA CRF neurons for 30 min during withdrawal (8 h) from chronic intermittent alcohol exposure on CeA Fos induction. The ANOVA revealed a significant effect of the optogenetic inhibition of CeA CRF neurons (*F*_3,2_ = 13.61, *p* < 0.01). The Newman Keuls *post hoc* test indicated that after inhibition (laser ON), the increase in Fos^+^ neurons that was normally observed during withdrawal (laser OFF; *p* < 0.01, laser OFF *vs*. naive) was completely prevented (*p* < 0.01, ON *vs*. OFF). Unilateral inhibition also partially affected the contralateral CeA, in which a significant decrease in Fos*^+^* neurons was observed on the contralateral side (*p* < 0.01, ON/contralateral *vs*. OFF; Fig. 4D).

### Dissection of CeA CRF pathways in alcohol drinking in dependent rats

#### Ventral bed nucleus of the stria terminalis

The two-way repeated-measures ANOVA, with time and laser as within-subjects factors, revealed significant effects of time (*F*_2,22_ = 9.18, *p* < 0.01) and laser (*F*_1,11_ = 29.72, *p* < 0.001) on alcohol drinking and a time × laser interaction (*F*_2,22_ = 5.31, *p* < 0.05). The Newman Keuls *post hoc* test showed a significant reduction of alcohol self-administration when the green laser was turned ON (*p* < 0.01, *vs*. laser OFF; Fig. 5A.a). A separate one-way ANOVA indicated that the animals exhibited escalation of alcohol intake compared with baseline pre-vapor exposure (*F*_6,66_ = 11.43, *p* < 0.0001). The Newman Keuls *post hoc* test indicated that rats showed escalation of alcohol drinking only on laser OFF days (*p* < 0.001), whereas the animals exhibited a level of responding that was similar to baseline pre-vapor exposure during optogenetic inhibition of CRF^CeA-BNST^ terminals (Fig. 5A.a). Immediately before the last two alcohol self-administration sessions, the rats were observed for withdrawal signs 8 h into withdrawal. As shown in Fig. 5A.b, the laser ON group exhibited significant decreases in abnormal gait (Mann-Whitney *U* = 6.00, *p* < 0.05) and body tremors (*U* = 10.50, *p* < 0.05). The sum of the five rating scores revealed a significant decrease in overall withdrawal severity (*U* = 1.5, *p* < 0.01; Fig. 5A.b, inset) after inhibition of CRF^CeA-BNST^ terminals. The *t*-test did not reveal an effect of the inhibition of CRF^CeA-BNST^ terminals on saccharin self-administration (*t*_11_ = 0.12, *p* > 0.05; Fig. 5A.c).

**Figure 5.**
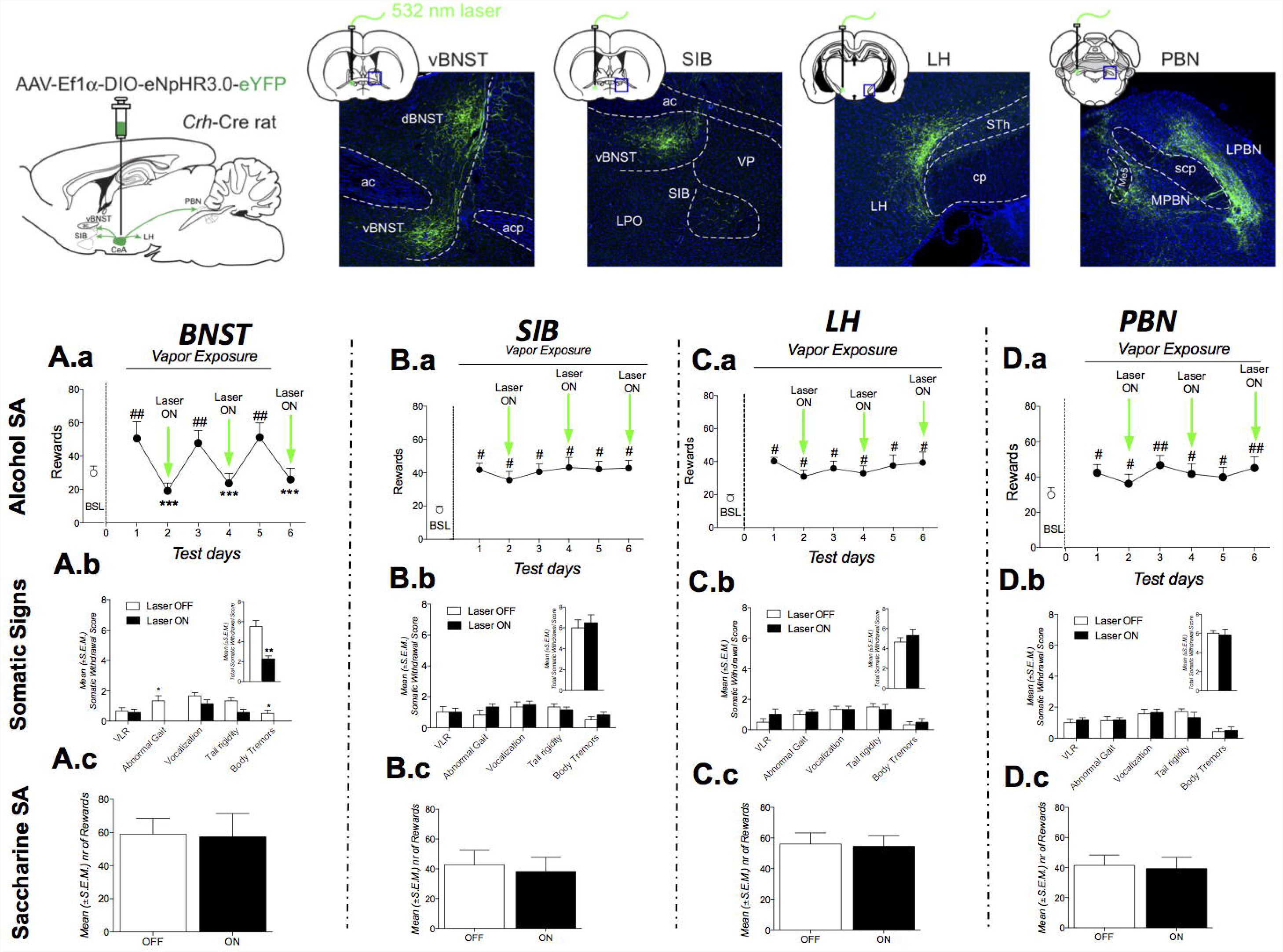
The schematic diagram shows representative images of the area of injection (CeA) and area of optogenetic inhibition (BNST, SI, LH, and PBN). (**A.a**) Time-course of the effect of unilateral optogenetic inactivation of CeA CRF BNST terminals. ^#^*p* < 0.05, *vs*. baseline (BSL); \*\**p* < 0.01, *vs*. laser OFF. (**A.b**) Effect of CRF^CeA-BNST^ terminal inhibition on somatic withdrawal signs. \**p* < 0.05, *vs*. laser OFF. (**A.c**) Effect of unilateral optogenetic inhibition of CRF^CeA-BNST^ terminals on saccharin self-administration. (**B.a**) Time-course of the effect of unilateral optogenetic inactivation of CRF^CeA-SI^ terminals. ^#^*p* < 0.05, *vs*. BSL. (**B.b**) Effect of CRF^CeA-SI^ terminal inhibition on somatic withdrawal signs. (**B.c**) Effect of unilateral optogenetic inhibition of CRF^CeA-SI^ terminals on saccharin self-administration. (**C.a**) Time-course of the effect of unilateral optogenetic inactivation of CRF^CeA-LH^ terminals. (**C.b**) Effect of CRF^CeA-LH^ terminal inhibition on somatic withdrawal signs. (**C.c**) Effect of unilateral optogenetic inhibition of CRF^CeA-LH^ terminals on saccharin self-administration. (**D.a**) Time-course of the effect of unilateral optogenetic inactivation of CRF^CeA-PBN^ terminals. ^#^*p* < 0.05, *vs*. BSL; \*\**p* < 0.01, *vs*. laser OFF. (**D.b**) Effect of CRF^CeA-PBN^ terminal inhibition on somatic withdrawal signs. \**p* < 0.05, *vs*. laser OFF. (**D.c**) Effect of unilateral optogenetic inhibition of CRF^CeA-PBN^ terminals on saccharin self-administration.

#### Substantia innominata

The two-way repeated-measures ANOVA, with time and laser as within-subjects factors, revealed a significant effect of time (*F*_2,10_ = 15.10; *p* < 0.001) but no effect of laser (*F*_1,5_ = 4.39, *p* > 0.05) and no time × laser interaction (*F*_2,10_ = 3.66, *p* > 0.05). A separate one-way ANOVA indicated that the animals exhibited escalation of alcohol intake compared with baseline pre-vapor exposure (*F*_6,30_ = 4.28, *p* < 0.01). The escalation of alcohol intake was unaffected by optogenetic inhibition of CRF^CeA-SI^ terminals, indicated by the Newman Keuls *post hoc* test. Operant responding for alcohol remained high on both the laser ON and OFF days (*p* < 0.05 and *p* < 0.01, respectively, *vs*. baseline; Fig. 5B.a). The optogenetic inhibition of CRF^CeA-SI^ terminals did not affect alcohol drinking (*t*_11_ = 0.49, *p* > 0.05). Immediately before the last two alcohol self-administration sessions, the rats were observed for withdrawal signs. As shown in Fig. 5B.b, inhibition of CRF^CeA-SI^ terminals did not affect overall withdrawal severity (*U* = 15.50, *p* > 0.05; Fig. 5B.b, inset). The *t*-test did not indicate an effect of inhibition of CRF^CeA-SI^ terminals on saccharin self-administration (*t*_5_ = 0.22, *p* > 0.05; Fig. 5B.c).

#### Lateral hypothalamus/parasubthalamic nucleus

The two-way repeated-measures ANOVA, with time and laser as within-subjects factors, revealed a significant effect of time (*F*_2,10_ = 8.87, *p* < 0.01) but no effect of laser (*F*_1,5_ = 0.78, *p* > 0.05) and not time × laser interaction (*F*_2,10_ = 0.16, *p* > 0.05; Fig. 5C.a). A separate one-way ANOVA indicated that the animals exhibited escalation of alcohol intake compared with baseline pre-vapor exposure (*F*_6,30_ = 4.28, *p* < 0.01). The escalation of alcohol intake was unaffected by optogenetic inhibition of CRF^CeA-LH/pSTN^ terminals, indicated by the Newman Keuls *post hoc* test. Operant responding for alcohol remained high on both the laser ON and OFF days (both *p* < 0.05, *vs*. baseline; Fig. 5C.a). Immediately before the last two alcohol self-administration sessions, rats were observed for withdrawal signs. As shown in Fig. 5C.b, inhibition of CRF^CeA-LH/pSTN^ terminals neurons did not affect the overall withdrawal severity (*U* = 14.00, *p* > 0.05; Fig. 5C.b, inset). The *t*-test did not indicate an effect of inhibition of CRF^CeA-LH/pSTN^ terminals on saccharin self-administration (*t*_5_ =1.024, *p* > 0.05; Fig. 5C.c).

#### Parabrachial nucleus

The two-way repeated-measures ANOVA, with time and laser as within-subjects factors, revealed a significant effect of time (*F*_2,22_ = 7.14, *p* < 0.01) but no effect of laser (*F*_1,11_ = 0.73, *p* > 0.05) and no time × laser interaction (*F*_2,22_ = 1.99, *p* > 0.05). A separate one-way ANOVA indicated that the animals exhibited escalation of alcohol intake compared with baseline pre-vapor exposure (*F*_6,66_ = 4.75, *p* < 0.001). The escalation of alcohol intake was unaffected by optogenetic inhibition of CRF^CeA-PBN^ terminals, indicated by the Newman Keuls *post hoc* tests (Fig. 5D.a). Immediately before the last two alcohol self-administration sessions, the rats were observed for withdrawal signs. As shown in Fig. 5D.b, inhibition of CRF^CeA-PBN^ terminals did not affect the overall withdrawal severity (*U* = 18.50, *p* > 0.05; Fig. 5D.b, inset). The *t*-test did not indicate an effect of inhibition of CRF^CeA-PBN^ terminals on saccharin self-administration (*t*_11_ = 0.77, *p* > 0.05; Fig. 5D.c).

## Discussion

The present study demonstrates that CRF neurons represent the majority of neurons that comprise the CeA neuronal ensemble that is recruited during withdrawal in alcohol-dependent rats. We further demonstrated that activation of CRF neurons in the lateral CeA (CeL) is required for the recruitment of a CeA neuronal ensemble that is located in the CeL, medial CeA (CeM), and capsular CeA (CeC). Optogenetic inhibition of CeA CRF neurons fully reversed the escalation of alcohol drinking and partially alleviated the somatic signs of withdrawal in dependent rats, without affecting water or saccharin self-administration. Finally, the selective optogenetic inhibition of CRF^CeA-BNST^ pathways but not CRF^CeA-SI^, CRF^CeA-LH/pSTN^, or CRF^CeA-PBN^ pathways recapitulated the behavioral effects of the inactivation of CeA CRF neurons.

Numerous studies have demonstrated that chronic intermittent exposure to alcohol vapor has robust predictive validity for alcoholism and construct validity for the neurobiological mechanisms of alcohol dependence (Heilig and Koob, 2007; Koob, 2009). Rats that are made dependent with chronic intermittent exposure to alcohol vapor (14 h on/10 h off) increase their alcohol drinking when tested during early and protracted withdrawal, with clinically relevant blood alcohol levels (150-250 mg%) and compulsive alcohol intake despite adverse consequences (Vendruscolo *et al*, 2012). Compulsive drug seeking that is associated with alcoholism can be derived from multiple neuroadaptations, but a key component involves the construct of negative reinforcement, defined as alcohol drinking to alleviate a negative emotional state (Koob, 2014). The recruitment of brain stress systems, such as CRF, in the extended amygdala has been hypothesized to play central role in generating the negative emotional state that drives such negative reinforcement.

We recently demonstrated that withdrawal from alcohol produces significant recruitment of Fos^+^ neurons in the CeA (de Guglielmo *et al*, 2016). We found that recruitment of the CeA neuronal ensemble is limited to a small subpopulation of neurons in the CeA, in which double labeling with Fos and NeuN revealed that only 7-8% of neurons (NeuN^+^) were Fos^+^ (Supplementary Fig. 1). No difference was observed between the different subregions of the CeA, although the lateral CeA (which contains the majority of CRF neurons; (Pomrenze *et al*, 2015) exhibited the largest increase in Fos^+^ neurons (Supplementary Fig. 1). Moreover, double Fos-CRF immunohistochemistry showed that the total number of CRF neurons did not change between groups (Fig. 1), but the number of Fos^+^/CRF^+^ neurons in the CeA dramatically increased during withdrawal from alcohol. Finally, neuronal phenotyping showed that the CeA withdrawal neuronal ensemble was mostly composed of both CRF^+^ and CRF^−^ neurons, with CRF^+^ neurons representing the majority (~80%) of the total number of Fos^+^ neurons.

We previously demonstrated that inactivation of the withdrawal neuronal ensemble in the CeA reverses the escalation of alcohol drinking in dependent rats and alleviates the somatic signs of withdrawal (de Guglielmo *et al*, 2016). However, unknown was whether the activation of CeA CRF neurons is a cause or consequence of recruitment of the CeA withdrawal neuronal ensemble. The results showed that the inactivation of CeA CRF neurons completely prevented recruitment of the CeA neuronal ensemble that includes both CRF^+^ and CRF^−^ neurons. These results are consistent with our previous work showing that chemogenetic stimulation of CeA CRF neurons recruited a population of CRF^+^ and CRF^−^ neurons in the CeL and CeM that was prevented (for CRF^−^ neurons) by a CRF_1_ receptor antagonist (Pomrenze *et al*, 2015). CeA CRF neurons are γ-aminobutyric acid (GABA)ergic, and the activation of CRF_1_ receptors in the CeL increases glutamate release locally (Silberman and Winder, 2013), suggesting that the recruitment of CRF^−^ neurons in the neuronal ensemble may be mediated by the activation of CRF_1_ receptors on glutamatergic terminals. Unilateral inhibition might also affect the contralateral CeA, suggesting the partial synchronization of bilateral CeAs. We next evaluated the effects of the optogenetic inhibition of CeA CRF neurons in animals that were trained to self-administer alcohol and were made dependent by chronic intermittent exposure to alcohol vapor. *Crh*-Cre rats were infused with AAV-DIO-NpHR-eYFP (eNpHR) or AAV-DIO-eYFP (eYFP) and bilaterally implanted with fiber optics in the CeA. The inactivation of CeA CRF neurons decreased alcohol drinking in dependent rats (Fig. 3). This effect was reversible, and inactivation reversed the level of drinking to pre-escalation levels before the induction of dependence. Altogether, we found that CeA CRF neurons are responsible for activation of the CeA neuronal ensemble and excessive alcohol drinking in dependent rats. Inactivation of CeA CRF neurons had no effect on saccharin self-administration or locomotor activity, and no effect of the laser was observed in the control group (AAV-DIO-eYFP; Fig. 3). These results suggest that the decrease in drinking that was observed with NpHR was specific to alcohol drinking in dependent rats and not attributable to nonspecific alterations of other behaviors. Although we did not observe nonspecific effects of 30 min inactivation using 532 nm/NpHR on behavioral measures, one concern may be potential neuroadaptations at the cellular level. To test this possibility, we tested the effects of only 5 min of the green laser in NpHR-injected rats because inhibition for 5 min using 532 nm/NpHR did not produce aberrant or nonspecific neuronal responses (Mahn *et al*, 2016). We did not observe any qualitative or quantitative differences between the 5 and 30 min exposure times. Indeed, the same effect size between NpHR and YFP rats was observed after 5 min compared with 30 min.

We then tested whether inactivation of the population of CeA CRF neurons during withdrawal would modulate the recruitment of Fos^+^ neurons in the CeA. We implanted *Crh*-Cre rats with unilateral fiber optics over the CeA and tested the effect of unilateral inhibition of CeA CRF neurons for 30 min during withdrawal (8 h) from chronic intermittent exposure to alcohol as in the previous experiment. Unilateral inhibition of CeA CRF neurons during withdrawal led to the same reduction of alcohol intake that was observed with bilateral inhibition. We also found that somatic withdrawal signs were significantly reduced. We recently found that CeA CRF neurons send dense projections to the BNST, SI, LH/pSTN, and PBN, but the role of each CRF projection in alcohol drinking was unknown.

Inactivation of the CRF^CeA-BNST^ pathway recapitulated the effects of silencing CeA CRF cell bodies, demonstrating that activation of the CRF^CeA-BNST^ projection is critical for excessive alcohol drinking in dependent rats (Fig. 5). The effect was specific to alcohol drinking and withdrawal since saccharin self-administration was unaffected.

The BNST is considered an integrative hub that connects several stress-related brain regions, including the basolateral amygdala, CeA, and PVN, and brain reward centers, such as the ventral tegmental area and nucleus accumbens (Brog *et al*, 1993; Georges and Aston-Jones, 2001, 2002; Rinker *et al*, 2016; Silverman *et al*, 1981). Importantly, the BNST is a critical neuropeptides, including CRF, innervate and modulate activity in the BNST (Kash *et al*, 2015). A substantial body of evidence supports the role of CRF signaling in the BNST during anxiety that is induced by withdrawal from drugs of abuse (Huang *et al*, 2010; Overstreet *et al*, 2003). These effects on withdrawal are thought to drive the effects of CRF in the BNST on stress-induced reinstatement of drug-seeking behavior (Erb *et al*, 2001; Erb and Stewart, 1999). The CeA sends dense CRF projections to the ventral and dorsal BNST (Pomrenze *et al*, 2015; Sakanaka *et al*, 1986). A clear role for BNST CRF signaling in stress-induced reinstatement has been reported, but less clear is the role of CRF signaling in the BNST in alcohol addiction. For example, although intra-CeA injections of CRF receptor antagonists after chronic intermittent alcohol exposure block chronic intermittent exposure-induced increases in alcohol self-administration (Funk *et al*, 2006; Varodayan *et al*, 2017), intra-BNST injections of the same antagonist post-chronic intermittent alcohol exposure have been shown to be ineffective (Funk *et al*, 2006). In the present study, we demonstrated a pivotal role for CRF^CeA-BNST^ projections in excessive alcohol drinking in dependent rats. The reduction of alcohol intake that was observed herein appears to be mediated by ventral BNST projections because they were the target of our fiber optics. We previously observed a lack of CRF cells from the ventral BNST in our *Crh*-Cre rats (Pomrenze *et al*, 2015).

No effect was observed in the CRF^CeA-SI^, CRF^CeA-LH/pSTN^, or CRF^CeA-PBN^. We identified a small portion of fibers that project from the CeA to the SI. Previous studies showed that acute alcohol does not affect Fos^+^ immunoreactivity in the SI (Herring *et al*, 2004), although chronic alcohol may lead to significant cellular loss in the SI in mice (Beracochea *et al*, 1987). The SI is a major attentional modulator of prefrontal cortex (PFC) function (Zaborszky *et al*, 1999). The CeA-SI-PFC circuit has not been studied in the context of drug addiction, but converging evidence suggests that it is involved in attentional processes, salience, and learning. We hypothesized that the CeA-SI-PFC pathway could promote alcohol drinking by increasing the salience of negative emotional states and facilitating learning to obtain relief from such emotional states through alcohol drinking. However, this hypothesis was not supported by the results because alcohol drinking and withdrawal signs were unaltered by inhibition of CRF^CeA-SI^ terminals. Another hypothesis that needs further testing is that CRF^CeA-SI^ terminals might be responsible for cognitive impairments that are caused by excessive alcohol drinking, given that PFC neuron recruitment during alcohol withdrawal predicts cognitive impairment. We observed a functional disconnection between the CeA and infralimbic/prelimbic cortex during abstinence from binge-like alcohol drinking (George *et al*, 2012).

The LH/pSTN is a major input to the medial habenula (MH)-interpeduncular nucleus (IPN) circuit, which has been recently shown to contribute to withdrawal-induced nicotine intake (Grieder *et al*, 2014; Zhao-Shea *et al*, 2015). Considering the critical role of the MH-IPN pathway in compulsive nicotine intake and behavioral signs of nicotine withdrawal, we expected that the optogenetic inhibition of CRF^CeA-LH/pSTN^ terminals would prevent excessive alcohol drinking and somatic signs of alcohol withdrawal. However, we did not observe such effects. Activation of this pathway during withdrawal was not an important factor in excessive drinking or the expression of somatic signs of withdrawal. An alternative hypothesis is that the CRF^CeA-LH/pSTN^ pathway is recruited during withdrawal but is specifically involved in the reinstatement of alcohol seeking in the absence of drinking. Indeed, the LH has been shown to be critical for the context-induced reinstatement of alcohol seeking (Moorman *et al*, 2016). Future studies of the context-induced reinstatement of alcohol seeking after the extinction of alcohol drinking are needed to test this hypothesis.

The largest and densest projection from CRF^CeA-PBN^ terminals was identified in the lateral PBN. The CeA receives nociception-related information via a direct monosynaptic pathway from the external part of the lateral PBN (Sarhan *et al*, 2005; Han et al., 2015 - PMID: 26186190), which is the predominant target of the ascending nociceptive spino-parabrachio-amygdaloid pathway (Jasmin *et al*, 1997; Todd, 2010). Previous studies showed that nociceptive stimuli increase neuronal activity (Hermanson and Blomqvist, 1996) and Fos immunoreactivity (Bester *et al*, 1997) in the PBN. The inhibition of CRF^CeA-PBN^ terminals did not affect alcohol drinking or the somatic signs of withdrawal, and one possibility is that the CRF^CeA-PBN^ projection mediates hyperalgesia during withdrawal from alcohol (de Guglielmo *et al*, 2017; Edwards *et al*, 2012). Further studies will be required to test this hypothesis.

A large body of evidence shows that CRF and GABA systems in the CeA play an important role in alcohol dependence (Koob, 2008; Menzaghi *et al*, 1994; Merlo Pich *et al*, 1995; Valdez and Koob, 2004). At the cellular level, acute alcohol enhances evoked GABA_A_ receptor-mediated inhibitory postsynaptic currents by increasing GABA release in rats (Roberto *et al*, 2003; Roberto *et al*, 2004) and mice (Nie *et al*, 2004). These effects are mediated by CRF and can be blocked by a CRF_1_ antagonist (Roberto *et al*, 2010). The effects on alcohol intake that were observed herein were specific to CRF neurons, but we cannot exclude the possibility that they might be mediated by the inactivation of other neurotransmitters, such as CRF, GABA, dynorphin, or somatostatin, considering the high degree of colocalization and overlap between these populations of neurons (Gilpin and Roberto, 2012; Kang-Park *et al*, 2015; Marchant *et al*, 2007; Pomrenze *et al*, 2015). Future studies should identify the neurotransmitters CeA CRF neurons release locally and in the BNST to mediate alcohol drinking and withdrawal signs in the dependent state.

In summary, the present study found that the activation of CeA CRF neurons was required for recruitment of the CeA neuronal ensemble that drives excessive alcohol drinking in dependent rats. Inactivation of the CRF^CeA-BNST^ pathway completely reversed excessive alcohol drinking during withdrawal and partially prevented the somatic signs of withdrawal. Inactivation of the CRF^CeA-SI^, CRF^CeA-LH/pSTN^, and CRF^CeA-PBN^ pathways did not impact alcohol drinking or the somatic signs of withdrawal. Future studies will be important to determine whether the CRF^CeA-SI^, CRF^CeA-LH/pSTN^, and CRF^CeA-PBN^ pathways are instead involved in mediating other aspects of alcohol dependence, including cognitive impairments, alcohol seeking, and hyperalgesia. These results suggest dissociable roles for CeA CRF pathways in addiction, and targeting the CRF CeA neuronal ensemble or CRF^CeA-BNST^ pathway may facilitate the development of novel therapeutic approaches for the treatment of alcohol use disorders.

## Acknowledgements

The authors thank Michael Arends for proofreading the manuscript.

## Footnotes

This study was supported by National Institutes of Health [Grant AA006420], [Grant AA020608], [Grant AA022977] (OG), [Grant AA13588], and [Grant AA017072] (ROM).

